# Nutrimental determinants of chronological aging and competitiveness in the *snf1Δ* Warburg model

**DOI:** 10.64898/2026.06.18.733183

**Authors:** Abigail Correa-Olivares, Amanda de la Caridad Lahera Champagne, Alma Delia Bertadillo-Jilote, Karla I. Lira-de Leon, David G. Garcia-Gutierrez, Gerardo M. Nava, Vanessa Sanchez-Quezada, Luis Alberto Madrigal-Perez

## Abstract

Cancer, one of the world’s leading causes of death, is characterized by a complex metabolic reprogramming that features the Warburg effect as one of its hallmarks. The Warburg effect involves increased glucose and amino acid metabolism, which promotes tumor proliferation and progression. Although cancer has historically been attributed to genetic mutations, recent studies suggest a possible metabolic origin. However, a key characteristic of cancer cells is their greater adaptability than normal cells, as evidenced by their resistance to chemotherapy, which stems from their high mutability. This underscores the need to examine the relationship between metabolic reprogramming and cancer development from both metabolic and evolutionary perspectives. In this context, *Saccharomyces cerevisiae snf1*Δ strain has emerged as an ideal cellular model for studying the Warburg effect. This study aimed to determine whether deletion of the *SNF1* gene in *S. cerevisiae* affects its chronological aging and competitiveness in a glucose and amino acid-dependent manner. Herein, we provide evidence that the *snf1*Δ strain changes the chronological aging depending on nutrimental condition, under low-nutrient levels shortens (0.1% glucose + 0.1x amino acids), and increases under high-nutrient levels (5% glucose + 3x amino acids). Competitiveness of the *snf1*Δ strain in co-cultivation with wild-type was also improved in 5% glucose + 3x amino acids, by approximately 2 Log10. These results indicate that *snf1*Δ strain aging and competitiveness are also sensitive to nutrimental status, as was observed in cancer cells.

## Introduction

Cancer, one of the leading causes of death worldwide, is associated with a series of metabolic alterations that promote the uncontrolled growth of tumor cells [1]. In 2020, according to data from the International Agency for Research on Cancer (IARC), an estimated 19.3 million new cases of cancer and almost 10 million deaths from this disease were diagnosed globally [2]. The study of cancer has been closely linked to DNA mutations as the main cause [3]. However, a possible metabolic origin for this condition has recently been considered, based on the Warburg effect [4, 5].

The canonical definition of the Warburg effect describes the accelerated glycolysis that occurs even under normoxic conditions, with the conversion of glucose to lactate [6]. Recently, the metabolic characterization of the Warburg effect has expanded beyond glucose metabolism to include amino acid metabolism [5]. Amino acids play an important role in maintaining the cellular fitness of tumor cells. For example, glutamine is coupled to ATP synthesis at the substrate level via glutaminolysis [7] and also serves to produce proline and maintain redox homeostasis in tumor cells [8].

Cancer cells use substrate-level phosphorylation to synthesize ATP, which is a pathway that produces fewer molecules of ATP per mole of oxidizable substrate than oxidative phosphorylation. However, in terms of ATP production rate, substrate-level phosphorylation is faster than oxidative phosphorylation [9]. Nonetheless, cells require more oxidizable substrates (e.g., glucose or glutamine) to maintain thermodynamically favorable ATP levels under conditions of substrate-level phosphorylation (Warburg effect). In this regard, it is expected that greater nutrient availability positively selects for cancer cells, as seen in obesity [10] and in gliomas under high-glucose and/or glutamine conditions [3, 7]. Coupling of energy synthesis to cellular proliferative capacity has gained attention. Interestingly, normal cells in division have been reported to exhibit a transient Warburg effect [11]. An association between substrate-level phosphorylation and growth rate has also been reported for *Saccharomyces cerevisiae* [12] and *Escherichia coli* [9]. Nonetheless, the point of view of the genetic origin of cancer has limited studies of the nutrimental impact on cancer cells.

A new model of the Warburg effect has emerged as an important tool to answer these questions. The *snf1*Δ strain of *S. cerevisiae* shared the metabolic and phenotypic features with the Warburg effect [13-15]. This yeast could help us answer how glucose and amino acid concentrations affect aging and competitiveness under the Warburg effect.

Therefore, this study aimed to analyze the chronological aging and cellular competitiveness of the *snf1*Δ strain of *S. cerevisiae* in relation to variations in glucose and amino acid levels, to explore the phenotype of this Warburg effect model. The results corroborated that, according to the theoretical Warburg effect phenotype, the *snf1*Δ strain is more responsible for nutrimental status in chronological aging and competitiveness. These results corroborated the sensitivity of the Warburg effect to nutrimental status explored in the *S. cerevisiae snf1*Δ strain.

## Material and methods

### Strains

The experiments were conducted using the genetic background of *S. cerevisiae* BY4742 (wild-type, *MaT*α; *his3*Δ*1; leu2*Δ*0; lys2*Δ*0; ura3*Δ*0)* and its mutant in the *SNF1* gene (*snf1*Δ, *MAT*α; *his3*Δ*1; leu2*Δ*0; lys2*Δ*0; ura3*Δ*0; YDR477w: kanMX4*), acquired from the EUROSCARF program (University of Frankfurt, Germany). These strains were maintained in YPD medium (2% glucose, 2% casein peptone, and 1% yeast extract). For the mutant strain, the medium was supplemented with the antibiotic geneticin (G-418, Sigma-Aldrich, St. Louis, MO, USA) at a final concentration of 200 μg/mL.

### Preparation of media and pre-inoculum

The pre-inoculum was prepared by streaking the strains onto Petri dishes containing 20 mL of YPD solid medium (2% glucose, 2% casein peptone, 2% bacteriological agar, and 1% yeast extract) supplemented with geneticin (200 μg/mL). These were incubated for 48 h at 30 °C to obtain isolated colonies. Then, one to three isolated colonies were transferred to a 10 mL test tube containing 5 mL of YPD liquid medium (2% glucose, 2% casein peptone, and 1% yeast extract) under aseptic conditions, and the test tubes were incubated for 24 h at 30°C with constant shaking at 250 rpm. For the mutant strains, geneticin was added at a final concentration of 200 μg/mL.

### Nutrimental experimental design

Different glucose concentrations were used to evaluate their effects on the phenotypes of wild-type and *snf1*Δ strains by supplementing the media with three levels: 0.1%, 1%, or 5%, representing low-, intermediate-, and high-glucose concentrations [16]. In the case of amino acids, they were supplemented, taking as reference 1 g per L of medium as 1x, for the low amino acid condition was supplemented at 10 times lower, 0.1 g per L of medium (0.1x), and the high amino acid concentration was 3 times the amino acid amount, 3 g per L of medium (3x).

### Bioreactor design: operating conditions

500 μL of pre-inoculum were inoculated into 50 mL Schott Duran flasks containing 10 mL of minimal SC medium (1% drop-out medium synthetic supplements without uracil, 0.5% ammonium sulfate, 0.2% dipotassium phosphate, 0.18% amino acid-free yeast nitrogen base, and 400 μg/mL uracil) supplemented with different concentrations of glucose, drop-out medium synthetic supplements without uracil (as the sole amino acid source), and geneticin (for the mutant strain).

For the agitation system, a 1 cm stirrer bar and ANZESER SH-2 magnetic stirrers were used. A 6 mm-diameter hole was drilled in the lids of the flasks requiring aeration, through which a 10 cm heat-resistant hose of the same diameter was inserted to connect a 0.2 μm AERVENT™ sterilizing-grade filter for gas exchange. Additionally, a syringe needle attached to a sterile 0.45 μm MICROLAB SCIENTIFIC PVDF syringe filter was inserted into the base of the lid. This mechanism was intended to allow for sample collection every 48 h using a sterile 5 mL syringe for the chronological aging assays (**Fig. 1**).

**Fig. 1.**
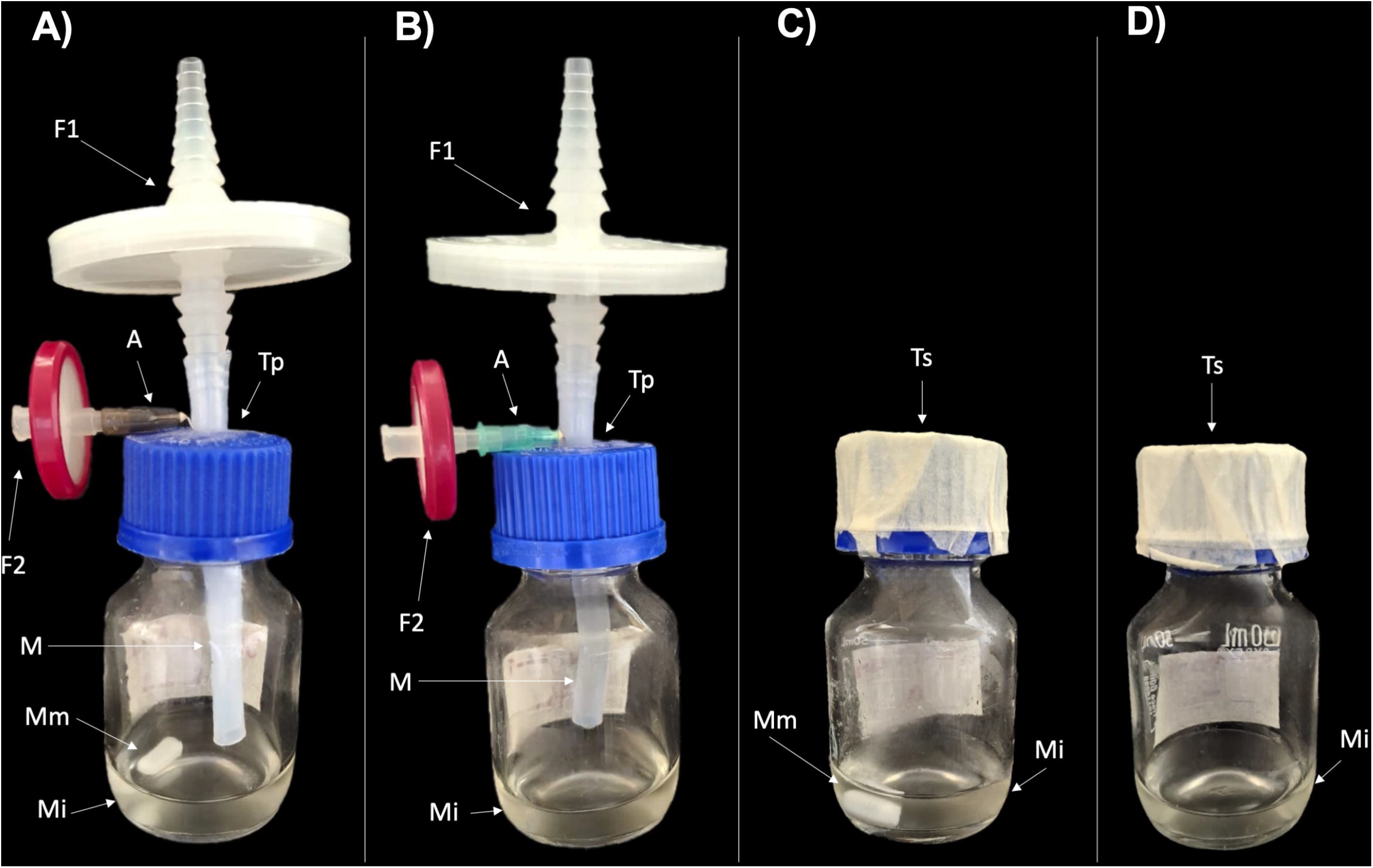
Bioreactor design. A) Bioreactor with agitation and aeration. B) Bioreactor with aeration. C) Bioreactor with agitation. D) Bioreactor without agitation and aeration. The letters indicate the different components of the bioreactors: 0.2 μm filter (F1), 0.45 μm filter (F2), sterile syringe needle (A), tubing (M), stirrer bar (Mm), inoculated SC medium (Mi), perforated lids (Tp), and sealed lids (Ts).

### Measurement of kinetic and operating parameters

Samples were taken after 24 h post-inoculation. Optical density was recorded at 600 nm (OD_600_) with a VELAB™ VE-5600UV UV-VIS spectrophotometer, using 200 μL aliquots diluted in 800 μL of distilled water (1:4) in 1.5 mL cuvettes. A blank solution of the uninoculated medium in distilled water, in the same proportion, was used. Other parameters considered were pH, biomass, and dissolved solids (°Brix). These values were recorded at both the beginning and the end of the experiment. To this end, 1 mL of the sample was centrifuged in 1.5 mL microtubes at 10000 × *g* for 3 minutes; the supernatant was transferred to another 1.5 mL microtube, and the measurements were recorded. The biomass was estimated by weighing the tube with the pellet on a VELAB™ VE-204EC analytical balance and subtracting the weight of the empty tube. The pH was determined using CIVEQ® reagent strips by immersing them in the supernatant for approximately 5 seconds. The dissolved solids (°Brix) in the supernatant were measured with a Milwaukee MA871 digital refractometer by placing ∼50 μL aliquots into the refractometer.

### Data analysis

The yield Yx/s was calculated to indicate the amount of biomass (x) generated per unit of substrate (s) consumed. This was done using the following equation:

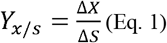

Where ΔX is the amount of biomass generated (g/mL) and ΔS is the amount of substrate consumed (°Brix).

On the other hand, the microbial growth curves were constructed using GraphPad Prism V. 11 software for Macintosh with the following equation:

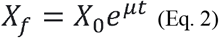

Where Xo is the initial cell population, Xf is the final cell population, μ is the specific growth rate, and t is time.

### Chronological aging

The chronological aging test was performed as described by Murakami, Kaeberlein [17] with some modifications. One to three colonies, each 1–1.5 mm in diameter, were inoculated into 50 mL Schott Duran flasks containing 10 mL of minimal liquid SC medium supplemented with varying concentrations of glucose, uracil-free drop-out medium (as the sole amino acid source), and geneticin (for the mutant strain). The flasks were incubated at 30°C with constant shaking at 80 rpm. The first sample was taken after the first 48 h of incubation and represented the first age point, while subsequent age points were collected over the next 2 weeks.

For each age point, growth kinetics were performed according to the method described by Olivares-Marin et al. [12]. 5 μL of each sample and 145 μL of SC minimal liquid medium were placed in a 96-well microplate, and 150 μL of SC minimal liquid medium was placed in a single well as the negative control.

The microplate was incubated for 24 h at 30 °C with constant shaking in a Varioskan Sky microplate reader (Thermo Scientific), which measured absorbance (at 600 nm) every 30 minutes. The data obtained were analyzed using GraphPad Prism V. 11 software for Macintosh.

For data analysis and growth curve generation, the data were normalized by subtracting the OD_600_ of the negative control from each column’s OD values. The growth kinetics were calculated using Equation 2. The doubling time (δ) was calculated for each well based on the growth kinetics at day 2, the age point. The equation used to calculate the doubling time was:

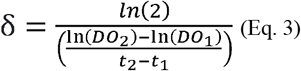

Where OD1 and OD2 represent successive OD measurements, and t1 and t2 are the times between measurements.

For each age point, the change in time (t) in the growth curves was calculated relative to the initial age point (day 2). An easy way to do this was to determine the difference in the time it takes for each well to reach an OD of 0.3 between the initial age point and each subsequent age point. The time it takes for a particular well to reach an OD of 0.3 can be calculated from the linear regression of ln(OD) on time, using the time between the two points marking OD = 0.3.

The survival rate at each age point was calculated to generate a survival curve. The initial age point (day 2) was defined to be 100% viable. For each successive age point, the survival percentage was calculated using the equation:

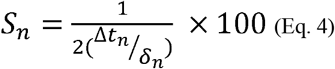

Where Sn is the survival percentage, Δtn is the change in time, and δn is the doubling time.

The results for both strains were plotted using the parameters established in Tables 3 and 4 to assess the influence of these changes in each case. The total area was calculated using area-under-the-curve analysis, and the results were plotted to confirm consistent behavior across all experiments.

### Competitiveness assay

For the cell competitiveness assay, 1 × 10□ cells of each strain were inoculated into 50 mL Schott Duran flasks containing 10 mL of minimal liquid SC medium supplemented with varying concentrations of glucose and synthetic Drop-out Medium without uracil (as the sole amino acid source): 0.1% glucose + 0.1x amino acids and 5% glucose + 3x amino acids. Each experiment also included a negative control to rule out contamination within the system. The flasks were incubated at 30°C with constant shaking at 80 rpm. The first sample was taken after the first 48 h of incubation (designated day 0), and subsequent samples were taken on days 10 and 20. Then, the wild-type and snf1Δ populations were determined by qPCR as follows. First, DNA isolation was performed according to the method of Edwards et al. [18] with some modifications.

Briefly, 2 mL of the homogenized sample was taken, and 1 mL was placed in a 1.5 mL microtube and centrifuged at 13000 × *g* for 10 min. The supernatant was discarded, and all traces of medium were removed from the pellet using a micropipette. The cells were resuspended in 1 mL of Edwards buffer (200 mM Tris pH 8.0, 200 mM NaCl, 25 mM EDTA, and 0.5% SDS). The samples were incubated for 10 min in a boiling water bath at 95 °C and centrifuged at 3000 × *g* for 10 min to remove cell debris. Subsequently, 400 µL of the supernatant was transferred to a new 1.5 mL microtube, and 400 µL of ice-cold isopropanol was added. To mix, the tube was gently inverted 5 to 10 times. The samples were incubated at room temperature for 10 min and centrifuged at 14000 × *g* for 10 min. The supernatant was decanted, and the DNA pellet was air-dried by inverting the microtube onto absorbent paper. Finally, the dried pellet was resuspended in 100 µL of sterile water for quantification and determine the quality using a microdrop plate in a Varioskan Sky spectrophotometer (Thermo Scientific). To confirm the integrity of the genetic material, electrophoresis was performed on a 1% agarose gel (CAS: 9012-36-6, Sigma-Aldrich) in 1X TAE buffer. Each well was loaded with 5 µL of DNA mixed with 5 µL of loading buffer (6X TriTrack DNA Loading Dye, SM0311 Thermo Scientific) and 3 µL of SYBR Green 1X (SYBR™ Green I nucleic acid gel stain, S7563 Thermo Scientific), using 5 µL of a 1 kb molecular weight marker (GeneRuler 1 kb DNA Ladder, SM0311 Thermo Scientific) as a reference. The gel was run at 100 V for 35 minutes with an approximate current of 70 mA. Finally, the gels were visualized on a photodocumentation system (GelSMART, DLAB) under UV light, and images of the bands were taken to verify DNA integrity and rule out degradation.

qPCR assays were performed on a CFX96 Touch™ real-time thermocycler (Bio-Rad, California, USA, Massachusetts, USA), using the commercial PowerUp™ SYBR® Green Master Mix for qPCR (Thermo Scientific, Massachusetts, USA). The qPCR assay mix consisted of 6 µL of PowerUp Master Mix, 0.48 µL of each qPCR primer (forward and reverse; **Table 1**), 0.6 µL of BSA (1:100), 1.44 µL of nuclease-free water, and 3 µL of DNA at a concentration of 5 ng/µL, for a final reaction volume of 12 µL per sample. Each sample was mounted in duplicate, as was each point on the standard curve. The PCR cycle programming and calibration curves were carried out as follows:

**Table 1.**
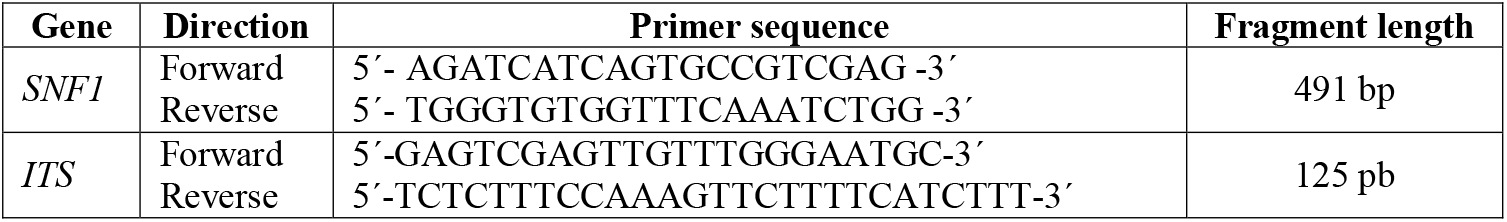
PCR primers used in this study.

Universal yeast (ITS):

UDG activation at 50°C for 2 min, initial denaturation at 95°C for 2 min, followed by 35 cycles of initial denaturation at 95°C for 30 sec, alignment at 60°C for 30 sec, and extension at 72°C for 30 sec, finally a cycle at 72°C for 3 min.

Efficiency: 102.6%

R^2^: 0.997

Slope: -3.26 Tm: 81°C

*SNF1*:

UDG activation at 50°C for 2 min, initial denaturation at 95°C for 2 min, followed by 35 cycles of initial denaturation at 95°C for 30 sec, alignment at 59.3°C for 30 sec, and extension at 72°C for 30 sec, finally a cycle at 72°C for 3 min.

Efficiency: 96.8%

R2: 0.993

Slope= -3.40

Tm: 82 °C – 81.50 °C

The DNA copy number was obtained using the formula:

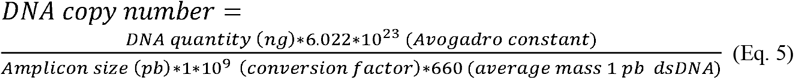

Subsequently, the relative abundance was obtained, and all data were converted to Log10 for data analysis.

### Statistical analysis

The statistical analysis was performed using GraphPad Prism V. 11 for Macintosh. All the experiments were carried out with at least three biological replicates. For mean comparisons of two groups, a t-test was made, and for three or more groups, a one-way ANOVA.

## Results

### Bioreactor design

Although the Warburg effect has traditionally been linked to mitochondrial dysfunction, mitochondrial respiration is highly active in Warburg effect cells, and maintaining this metabolic activity appears important [19]. Traditional methodologies for chronological aging assays use sealed test tubes with screw caps to house the cultures, which restricts gas exchange and yeast metabolism, inducing hypoxic conditions that limit mitochondrial respiration and have pleiotropic effects on metabolism. Therefore, to improve oxygenation and determine the ideal conditions for *snf1*Δ strain growth, different aeration and agitation scenarios were evaluated to assess their impact on gas exchange and oxygen availability in the medium. It was decided to manufacture bioreactors capable of maintaining constant gas exchange, agitation, and temperature. Gas exchange in the bioreactor is a particularly relevant factor in our study model because oxygen is a key component. Thus, specifying a bioreactor design that avoids hypoxia and maintains optimal growth was a fundamental part of the study.

For the wild-type strain, significant differences in growth were observed 24 h post-inoculation, dependent on aeration and agitation conditions (**Fig. 2**). The bioreactor combining agitation and aeration showed the greatest growth compared to the other bioreactors configuration (**Fig. 2**). On the other hand, the lowest growth was observed in the bioreactor evaluated only with aeration; although there were no significant differences between this bioreactor and the bioreactor without agitation or aeration, the growth of the latter was significantly lower compared to the bioreactor with agitation (**Fig. 2**). However, despite similar behavior between agitation without aeration and aeration without agitation, the significance was greater (**Fig. 2**).

**Fig. 2.**
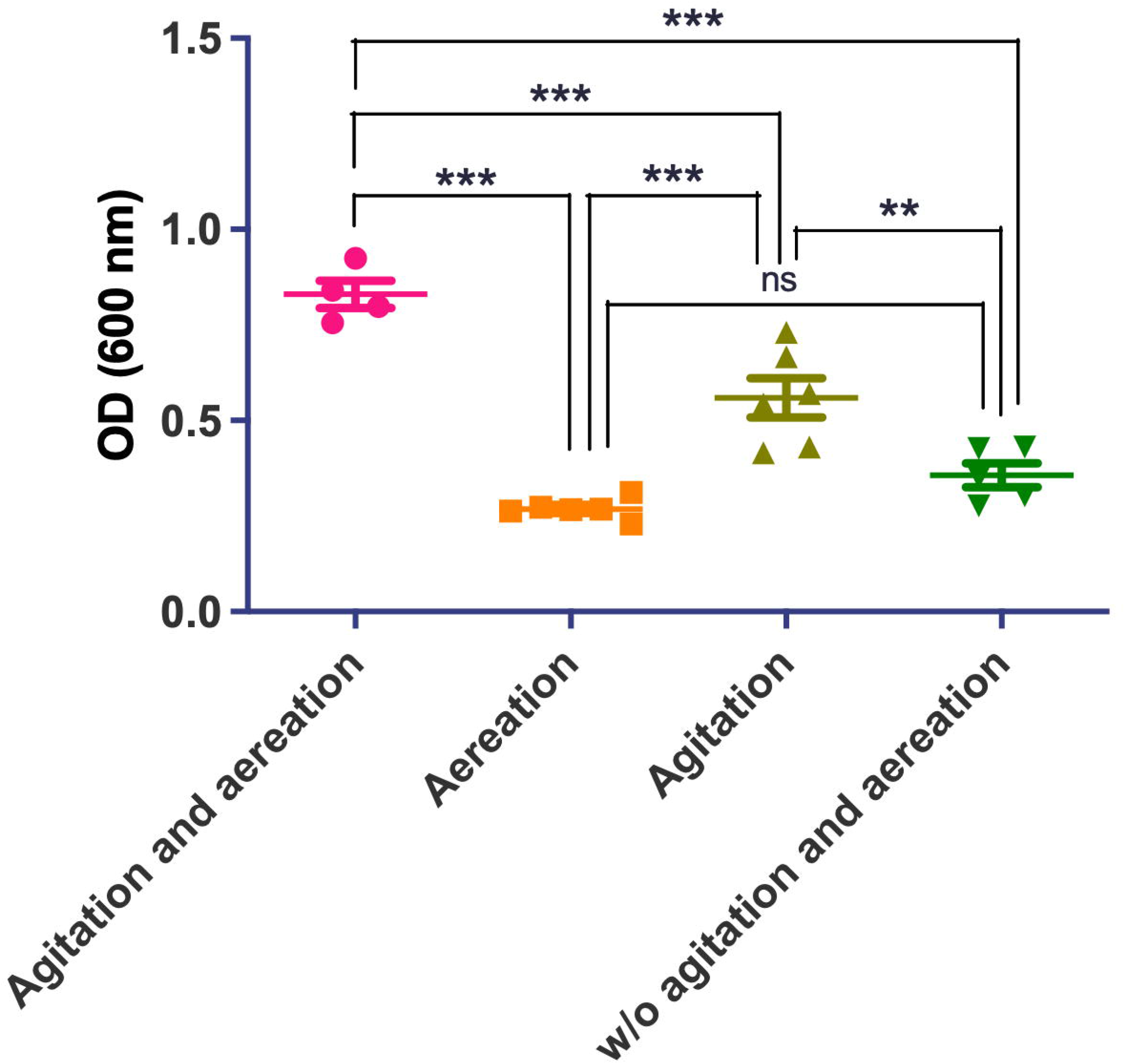
Growth of the wild-type strain at 24 h post-inoculation in the different bioreactor systems. Values represent the mean ± standard deviation of four to six independent experiments, including mean values from three technical replicates. Differences between means were analyzed by one-way ANOVA followed by Tukey’s test (ns = not significant, \**P* ≤ 0.05, \*\**P* ≤ 0.01, \*\*\**P* ≤ 0.001).

Regarding pH, no changes were generally observed between the final and initial measurements (**Table 2**). On the other hand, the highest biomass yield relative to the substrate was obtained with the bioreactor combining aeration and agitation (0.037 ± 0.011), followed by agitation alone (0.0314 ± 0.013), without aeration and agitation (0.0181 ± 0.031), and with aeration alone (0.0054 ± 0.013) (**Table 2**).

**Table 2.**
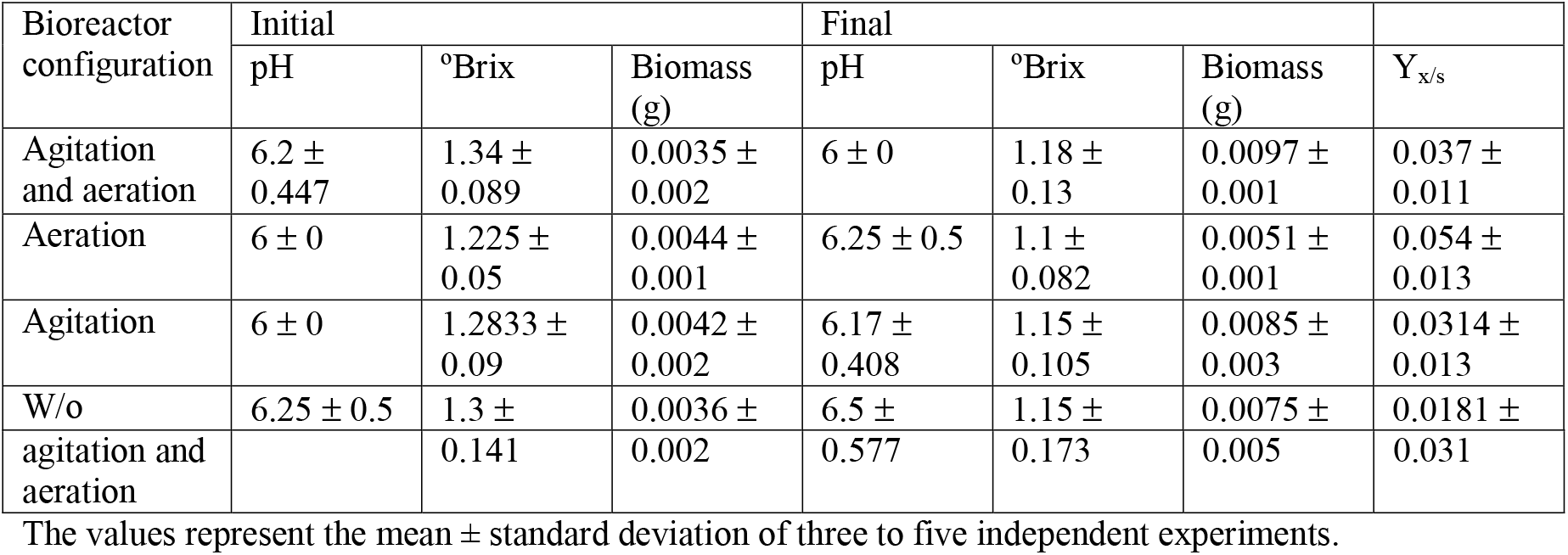
Kinetic and operating parameters for the wild-type strain under different bioreactor systems.

**Table 3.** Kinetic and operating parameters for the *snf1*Δ strain under different bioreactor systems.

| Bioreactor configuration | Initial | | | Final | | | $Y_{x/s}$ |
| --- | --- | --- | --- | --- | --- | --- | --- |
| | pH | $^{\circ}$Brix | Biomass (g) | pH | $^{\circ}$Brix | Biomass (g) | |
| Agitation and aeration | 6.5 ± 0.577 | 1.3 ± 0.082 | 0.0049 ± 0.001 | 6.25 ± 0.5 | 1.1 ± 0.141 | 0.0097 ± 0.001 | 0.0293 ± 0.013 |
| Aeration | 6 ± 0 | 1.2667 ± 0.058 | 0.0035 ± 0.001 | 6 ± 0 | 1.2667 ± 0.058 | 0.0053 ± 0.001 | Nd |
| Agitation | 6.2 ± 0.447 | 1.28 ± 0.045 | 0.0054 ± 0.002 | 6.2 ± 0.447 | 1.12 ± 0.045 | 0.0118 ± 0.003 | 0.0419 ± 0.016 |
| W/o agitation and aeration | 6.25 ± 0.5 | 1.325 ± 0.05 | 0.0053 ± 0.003 | 6.25 ± 0.5 | 1.15 ± 0.1 | 0.0031 ± 0.001 | -0.023 ± 0.031 |

The values represent the mean ± standard deviation of three to five independent experiments, nd: not determined.

The growth of the *snf1*Δ strain showed notable differences across the bioreactor configurations evaluated. Both the bioreactors with agitation and aeration and the bioreactors with agitation showed the greatest growth among all conditions (OD ∼1), but without a significant difference between them; interestingly, the lowest growth (OD ∼0.3) was observed in the bioreactor with aeration as well as the bioreactor without aeration or agitation, with no significant differences between these, but significantly lower compared to the previous conditions (**Fig. 3**). As with the parental strain, no changes were generally observed between the final and initial pH measurements for the mutant strain (**Table 3**). On the other hand, the highest biomass yield relative to the substrate was obtained with the agitated bioreactor (0.0419 ± 0.016), followed by the conditions with agitation and aeration (0.0293 ± 0.013) (**Table 3**). It should be noted that, for the aerated bioreactor, even with biomass production that could be comparable to that of the system without aeration and agitation, the yield could not be calculated because there was no change in °Brix (**Table 3**).

**Fig. 3.**
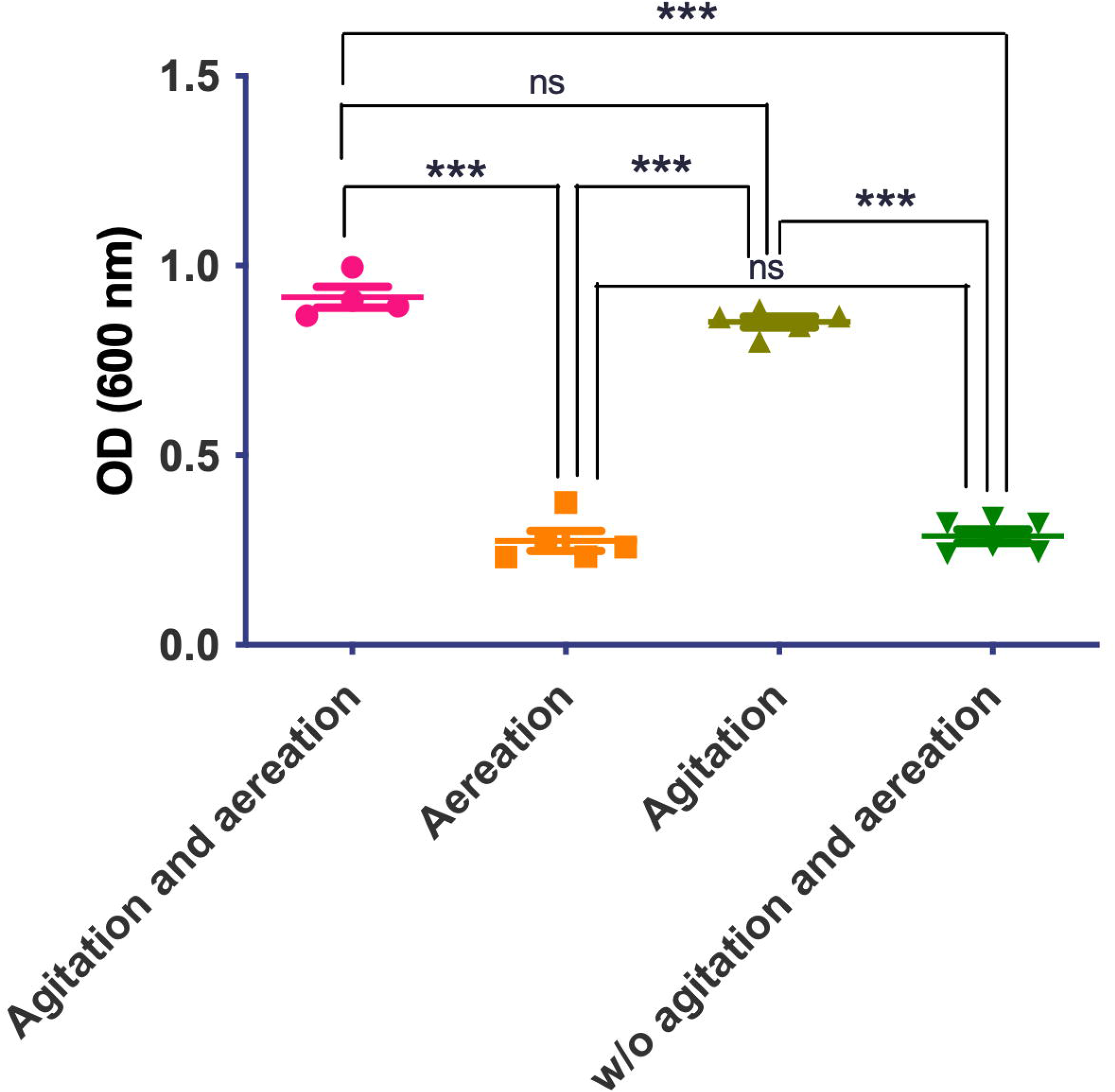
Effect of different bioreactor configurations on growth of the *snf1*Δ strain 24 h post-inoculation. Values represent the mean ± standard deviation of four to six independent experiments, including mean values from three technical replicates. Differences between means were analyzed by one-way ANOVA followed by Tukey’s test (ns = not significant, \**P* ≤ 0.05, \*\**P* ≤ 0.01, \*\*\**P* ≤ 0.001).

When comparing the WT and *snf1*Δ strains under different aeration and agitation conditions, notable differences in their behavior were observed. When a bioreactor with aeration and agitation was used, maximum growth was achieved in both strains, suggesting that this setup promotes better oxygen dispersion and more uniform nutrient distribution. However, when analyzing the effect of each factor separately, different responses were observed between strains: in the parental strain, both agitation and aeration contributed significantly to growth (**Fig. 2**), while in the mutant strain, growth was comparable under agitated conditions with and without aeration (**Fig. 3**).

Therefore, to continue the study of chronological aging, bioreactors equipped with a system that implements both agitation and aeration were chosen, as this would support microbial growth and performance and enable continuous gas exchange, as initially desired.

### Chronological aging

To assess how glucose and amino acid levels impact the Warburg effect model (*snf1*Δ), the growth of both strains was analyzed individually over 14 days using different media with varying levels of glucose (0.1%, 1%, and 5%) and amino acids (drop-out mix at 0.1x, 1x, and 3x). Studying the aging of both the parental and mutant strains under these conditions allowed us to identify the role of nutrient availability in the medium.

Treatments under which no significant variance in specific growth rate was detected included combinations of 0.1% and 1% glucose with 3x amino acids, as well as 5% glucose + 0.1x and 1x amino acids, and 0.1% of both glucose and amino acids (**Fig. 4**). On the other hand, the mutant strain showed a significantly higher growth rate than the wild-type strain under medium and low amino acid levels (1x and 0.1x, respectively) when supplemented with 1% glucose and 0.1% glucose + 1x amino acids (**Fig. 4**). Notably, the only treatment in which a significantly greater difference was observed was with 5% glucose + 3x amino acids, where the growth rate of the *snf1*Δ strain was higher than that of the wild-type strain (**Fig. 4**).

**Fig. 4.**
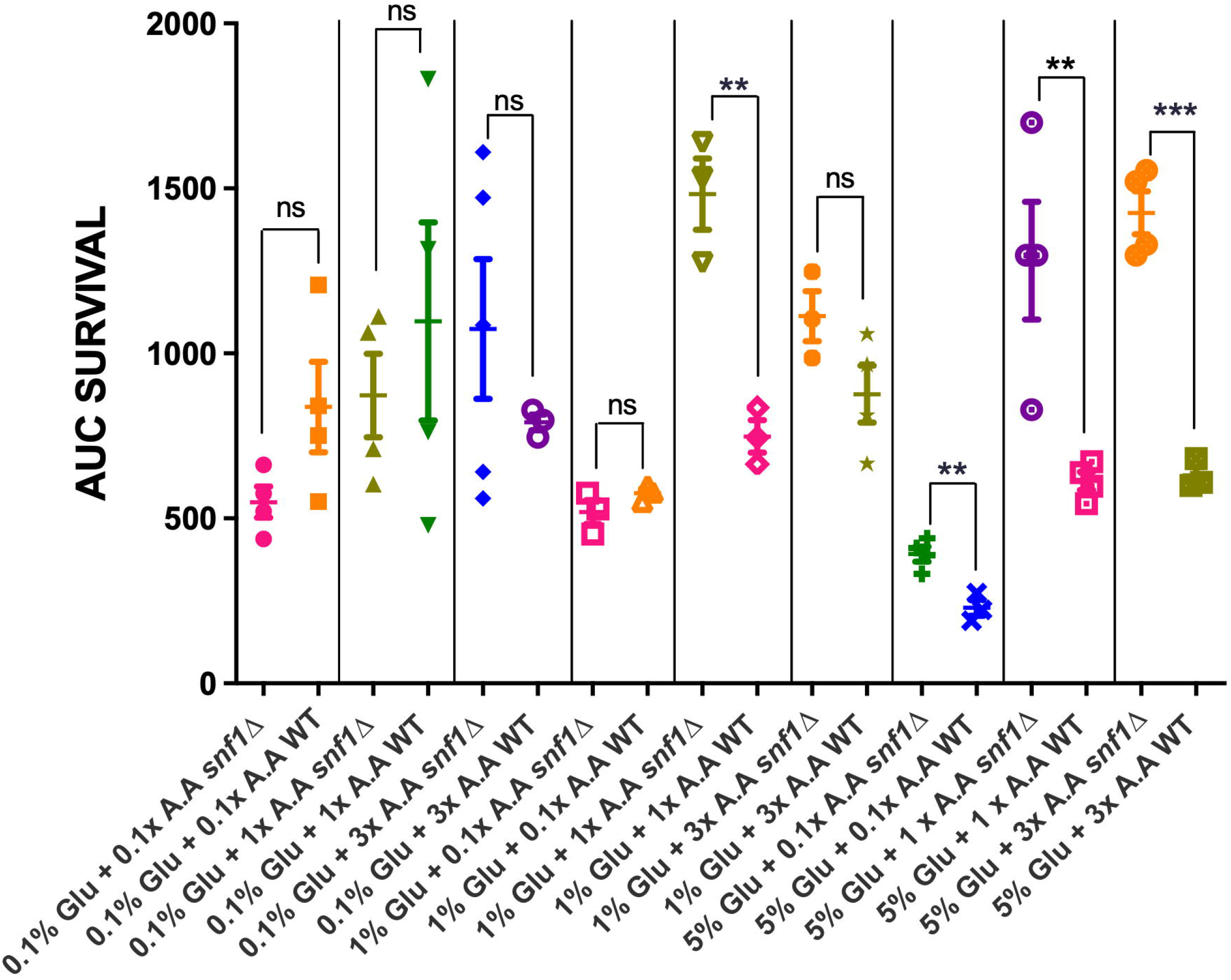
Influence of supplementation with different concentrations of glucose (Glu; 0.1%, 1%, 5%) and amino acids (AA; 0.1x, 1x, 3x) on chronological aging. Chronological aging of the wild-type and the *snf1*Δ strains was computed to obtain the area under the curves (AUC) to obtain an absolute value of chronological aging. Values represent the mean ± standard deviation of three to five independent experiments, including mean values from three replicates. Differences between means were analyzed using a two-tailed unpaired t-test (ns = not significant, \**P* ≤ 0.05, \*\**P* ≤ 0.01, \*\*\**P* ≤ 0.001).

In general, the behavior of the strains varied according to the nutritional conditions. Under glucose and amino acid restrictions, the wild-type strain survived better than the mutant, whereas at high concentrations of glucose and amino acids, the *snf1*Δ strain survived better than the parental strain. On the other hand, as nutrient availability in the medium increased, the aging of the mutant strain slowed compared to the wild-type strain (**Fig. 5A-I**). This behavior was particularly noticeable in the treatment supplemented with 5% glucose (**Fig. 5G-I**) and especially in the 3x amino acid supplementation (**Fig. 5I**). Specifically, the cell viability percentage of the wild-type strain, in contrast to the *snf1*Δ strain, was lower over time under caloric restriction (0.1% glucose; **Fig. 5 A-C**) combined with low (0.1x; **Fig. 5A**) and medium (1x; **Fig. 5B**) amino acid levels, contrary to what occurred with high amino acid levels (3x; **Fig. 5C**). This behavior persisted under medium glucose conditions (1%; **Fig. 5 D-F**) with 0.1x amino acids (**Fig. 5D**) but reversed when amino acid levels were increased (**Fig. 5E and 5F**). Interestingly, when exposed to high glucose levels (5%), the cell viability percentage of the snf1Δ strain was visibly higher than that of the parental strain in combination with all the amino acid levels evaluated (0.1x, 1x, and 3x; **Fig. 5 G-I**).

**Fig. 5.**
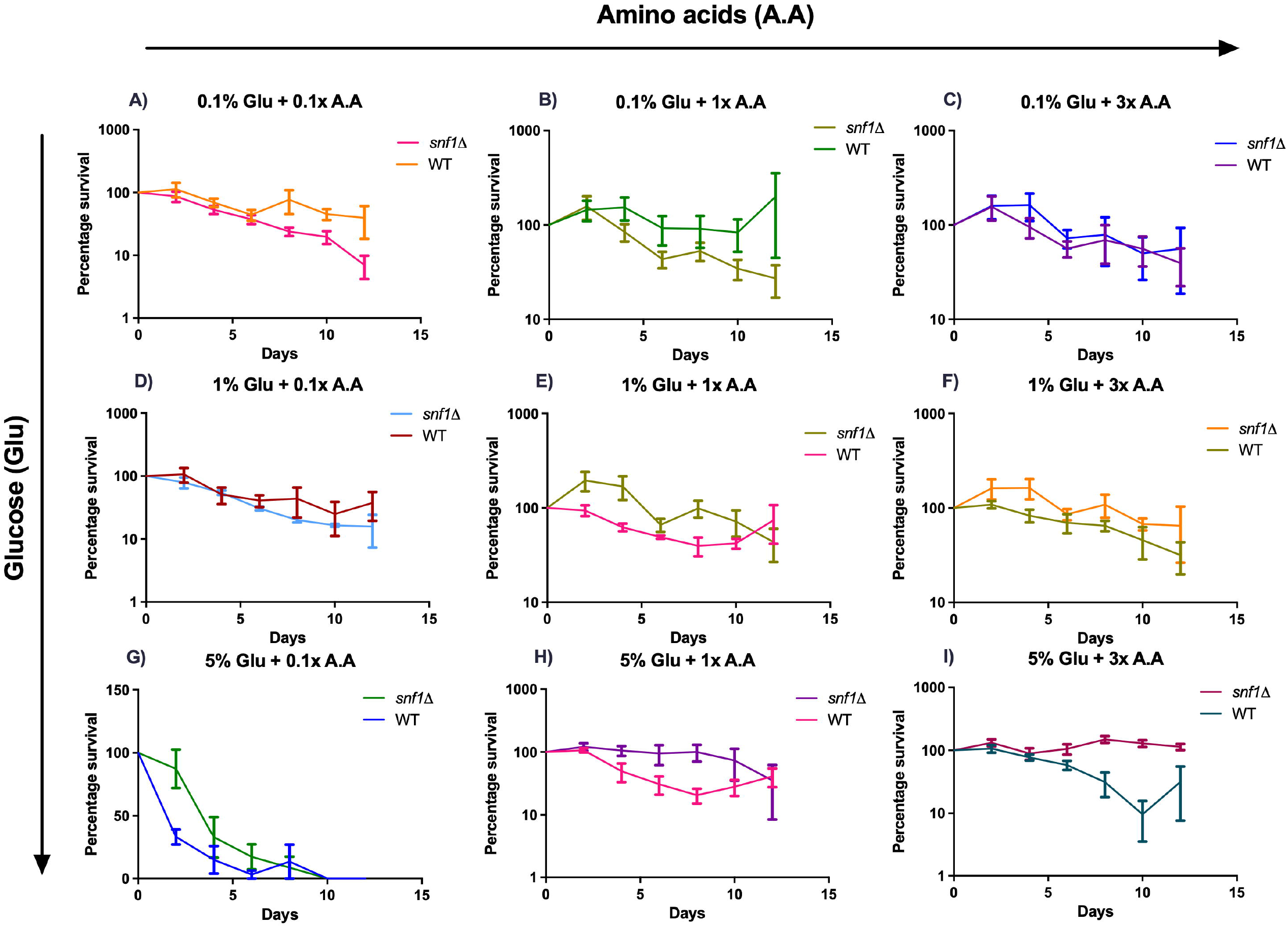
Chronological aging of the wild-type and the *snf1*Δ strains under different nutrimental conditions. Effect of supplementation with different concentrations of glucose (Glu; 0.1%, 1%, 5%) and amino acids (AA; 0.1x, 1x, 3x) on the percentage of cell viability of the wild-type and *snf1*Δ strains over 14 days. A) 0.1% glucose + 0.1x amino acids. B) 0.1% glucose + 1x amino acids. C) 0.1% glucose + 3x amino acids. D) 1% glucose + 0.1x amino acids. E) 1% glucose + 1x amino acids. F) 1% glucose + 3x amino acids. G) 5% glucose + 0.1x amino acids. H) 5% glucose + 1x amino acids. I) 5% glucose + 3x amino acids. The values represent the mean ± standard deviation of three to five independent experiments, including mean values from three technical repetitions.

Additionally, outgrowth points were detected in the viability curves of the mutant and wild-type strains, where cell viability increased again after a point of decline. Generally, these points were more pronounced under conditions where strain survival was compromised: for *snf1*Δ, they were observed under restrictive nutrient levels such as 0.1% glucose and 0.1x or 3x amino acids (**Figs. 5A and 5C**), or 1% glucose and 1x or 3x amino acids (**Figs. 5E and 5F**); for the wild-type strain, this was most noticeable in the treatments with 5% glucose and 1x and 3x amino acids (**Figs. 5H and 5I**). Furthermore, the only condition in which both strains reached total decline was in the treatment supplemented with 5% glucose and 0.1x amino acids (**Fig. 5G**).

Area under the curve (AUC) analysis revealed no significant differences between the wild-type and *snf1*Δ strains for any of the treatments, except for 1% glucose + 1x amino acids and 5% glucose supplemented with low (0.1x) and medium (1x) levels of amino acids, with a significantly higher AUC for the mutant strain compared to the wild-type. Likewise, the only treatment in which the AUC of *snf1*Δ was significantly higher than that of the parental strain was 5% glucose with 3x amino acids. Thus, under energetically favorable conditions (both glycolytic and amino acid-based), the area under the curve of cell viability percentage for the mutant strain significantly exceeded that of the wild-type.

### Competitiveness assays

An important phenotype of cancer cells is increased cell growth and competitiveness compared with normal cells. As *snf1*Δ is proposed as a Warburg effect model and according to the hypothesis that this metabolic phenomenon is key in cancer cells, we study whether *snf1*Δ cells prevail over wild-type in the two opposite nutrimental conditions that we assay. Chronological aging co-cultures with wild-type and *snf1*Δ strains were made to evaluate the competitiveness under two nutrimental conditions: 0.1% glucose + 0.1x amino acids, and 5% glucose + 3x amino acids. To evaluate the variation of wild-type and *snf1*Δ yeast populations, we use absolute DNA quantification using qPCR assays with oligonucleotides for amplification of *SNF1* and 16S rRNA genes. Since *snf1*Δ cells have the *SNF1* ORF interrupted, we obtain only the *SNF1* amplification corresponding to wild-type cells. To quantify the number of *snf1*Δ cells, the total yeast population was quantified by 16S rRNA and subtracted from the *SNF1* quantification. As expected, we found that the *snf1*Δ strain showed more fitness to prevail in high nutrimental conditions (5% glucose + 3x amino acids) than the wild-type strain (**Fig. 6B**). At this condition, the snf1Δ population increased 1.653 (Log10) and wild-type strain decrease 0.31 (Log10) (**Fig. 6B**). At low nutrimental conditions, we did not observed changes in both strains populations (**Fig. 6A**). This data corroborated that snf1Δ improves competitiveness in high nutrimental conditions, as expected by the Warburg effect model.

**Fig. 6.**
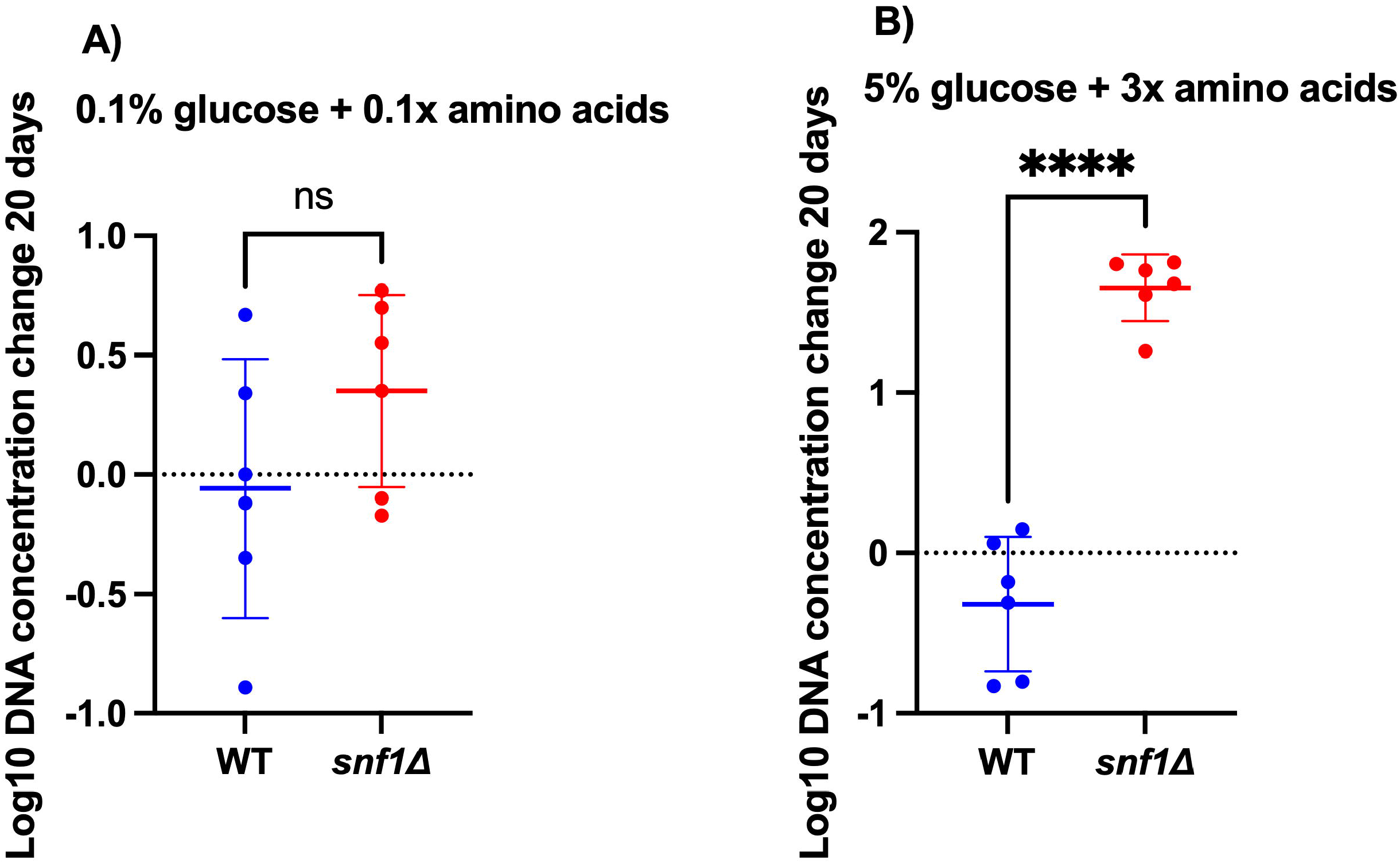
Competitiveness assay of co-culture of the wild-type and the *snf1*Δ strains under different nutrimental conditions. DNA quantification data were normalized using Log10 to obtain small numerical values. The values represent the mean ± standard deviation of six independent experiments. Differences between means were analyzed using a two-tailed unpaired t-test (ns = not significant, \*\*\**P* ≤ 0.001).

## Discussion

Although cancer research has historically focused on genetic alterations, a new approach has recently emerged, suggesting a possible metabolic origin for this disease, based on the Warburg effect [20]. This phenomenon indicates an increase in glycolytic flux directed toward fermentative pathways even under normoxic conditions [21]. This metabolic reprogramming enables tumor cells to sustain rapid cell proliferation by increasing the uptake of nutrients such as glucose and glutamine [4, 5]. However, current study models are based on mammalian cells, resulting in genetically variable and highly complex systems. In line with this, it has been proposed that the *S. cerevisiae* strain with a deletion of the *SNF1* gene, an AMPK homolog, exhibits metabolic characteristics reminiscent of the Warburg effect, making it a promising and unprecedented subject of study. This study aimed to determine whether the *snf1*Δ strain responds differentially to nutrimental conditions as the Warburg effect foresees in comparison to the wild-type strain, increasing its competitiveness. In this paper, there are two key findings: the *snf1*Δ strain shows slow chronological aging under high-nutrient conditions, and this improves its competitiveness compared to the wild-type strain also at high nutrimental condition. The results obtained in this study demonstrate that the behavior of the wild-type and *snf1*Δ strains of *S. cerevisiae* is strongly influenced by environmental conditions, particularly oxygen availability, agitation, and nutrient concentration.

Using a system that allowed us to control temperature, aeration, and oxygenation (**Fig. 1**), both strains showed maximum growth (**Fig. 2 and Fig. 3**). However, the *snf1*Δ strain showed greater sensitivity to the lack of agitation (**Fig. 3**), while the WT strain benefited from both agitation and aeration (**Fig. 2**). Therefore, agitation might be a more sensitive condition for the mutant strain, since under caloric (0.1%) and amino acid (0.1x) restrictions, it prioritizes uptake of these nutrients. On the other hand, the parental strain, lacking these characteristics in common with the Warburg effect, prioritizes oxygen uptake. When analyzing chronological aging, a response dependent on glucose and amino acid levels was observed. Under nutrient restriction conditions (0.1% glucose and 0.1x or 1x amino acids), the WT strain showed greater viability (**Fig. 5A and 5B**). Conversely, the mutant strain maintained greater viability over time compared to the parental strain under conditions of high nutrient availability (5% glucose and 3x amino acids) (**Fig. 5I**). This pattern, in which the prevalence of the mutated strain over the wild-type strain increased with increasing amino acid concentration and glucose level, was evidenced in the survival percentage graphs over time (**Figs. 5C and 5E-I**) and in the AUC graph of these same data (**Fig. 4**). Although no significant difference was observed in most treatments, there were conditions under which the *snf1*Δ strain outperformed the wild-type strain (1% glucose + 1x amino acids, and 5% glucose + 0.1x or 3x amino acids) (**Fig. 4**). Furthermore, it is noteworthy that the aging of the mutated strain slowed compared to the parental strain under the 5% glucose treatments with 1x and 3x amino acids (**Figs. 5H and 5I**). This behavior may be related to phenomena observed in tumor cells. For example, overexpression of glycolytic enzymes has been associated with various types of cancer, such as lung carcinoma [22]. Likewise, elevated levels of certain enzymes (transaminases BCAT1 and BCAT2) have been reported during the progression of chronic myeloid leukemia (CML) and acute myeloid leukemia (AML) [23]. Conversely, restriction of these nutrients has very marked negative consequences for cells. In prostate cancer, a lack of amino acids such as tyrosine and glutamine induces apoptosis and alters survival signaling pathways (e.g., Akt, Raf) and mitochondrial function [24]. Similarly, the viability and clonogenic capacity of HCT116 colorectal cancer cells were affected by a combination of glucose restriction, autophagy inhibition, and chemotherapy, with low-glucose supplementation further enhancing these effects [25].

In contrast, cancer cells have demonstrated extensive plasticity and adaptability to their environment, enabling them to sustain themselves through alternative pathways when necessary. When glucose or glutamine levels, the preferred nutrients of tumor cells, are restricted, there is a tendency to increase the consumption of other nutrients [26]. For example, in glioblastoma cells (GBM), activating serine synthesis from glucose enables them to maintain their proliferative capacity despite serine/glycine limitation in the brain microenvironment, making them especially aggressive and resistant [27]. In addition to what was found in this study, the viability curves (**Fig. 5**) show outgrowth phenomena, where, after an initial drop in viability, some cells manage to reactivate and proliferate; specifically, changes were observed in the nutritional conditions that affected their viability (for *snf1*Δ: 0.1% glucose + 0.1x and 3x amino acids (**Fig. 5A and 5C**), or 1% glucose + 1x or 3x amino acids (**Fig. 5E and 5F**), and for the WT strain: 5% glucose + 1x and 3x amino acids (**Fig. 5H and 5I**). This behavior shown by the WT and *snf1*Δ strains suggests that the cells can adapt to environmental conditions, a phenomenon reported in cellular aging. In this context, it has been shown that superoxide can act as a mediator of an “altruistic aging” program in *S. cerevisiae* [28]; this proposes that cells have the capacity to age and die in a programmed manner in order to release nutrients into the environment that allow for the survival and proliferation of a better-adapted subpopulation. This cellular dynamic has been reported in tumor cells; for example, pancreatic adenocarcinoma (PDA) cells subjected to low glucose (0.5 mM) and glutamine (0.1 mM) for several weeks can adapt by reactivating proliferation and increasing their carcinogenic capacity [26].

This type of evidence has enabled the recent development of therapeutic strategies such as the “Press Pulse” therapy, which combines the simultaneous restriction of glucose and glutamine with the induction of ketosis [4]. It has been shown that reducing glucose and increasing ketone body production decrease tumor progression, so that when the glucose-to-ketone ratio is low, cancer cells lack sufficient energy for their development [29]. For example, 6-diazo-5-oxo-L-norleucine (DON), a glutamine antagonist, was used in conjunction with a ketogenic diet with reduced caloric intake to treat advanced-stage glioblastoma (GBM) in two syngeneic murine orthotopic models. This approach reversed disease symptoms, eliminated tumor cells, and improved overall survival in the specimens [29]. This type of treatment has been used in real clinical cases [30].

In particular, the case of a patient with advanced IDH1-mutant GBM has been documented. This patient was managed without standard treatment and with a ketogenic diet, which mitigated tumor growth until a reductive surgical resection was performed. Subsequently, the patient showed prolonged survival of 80 months without major complications, suggesting a possible metabolic synergy between a low glucose/ketone ratio and the mutation, as they simultaneously affect glycolysis and glutaminolysis, which could contribute to long-term tumor control [30]. Similarly, the exclusive use of ketogenic metabolic therapy against a canine mast cell tumor, in conjunction with a 40% caloric restriction in the diet, has been reported to gradually reduce the tumor until its complete disappearance over several months, with no evidence of recurrence at the time of the specimen’s death years later [31].

Finally, the increase in competitiveness of the *snf1*Δ strain over the wild-type (**Fig. 6**) is in accordance with the main risk factors of cancer that are related to nutrimental status [32] and energy expenditure that causes a restriction of nutrients, preventing cancer for this reason [33]. This experiment is the proof of how the *snf1*Δ Warburg effect model could facilitate the study of important nutrimental status poorly addressed in mammalian cell lines.

In conclusion, all the findings obtained throughout the study help us to consolidate the theory that the deletion of the *SNF1* gene in *S. cerevisiae* can induce a Warburg-type metabolism, and that its behavior is favored for the availability of nutrients in its environment.

## Acknowledgments

This study was funded by the Facultad de Química of Universidad Autónoma de Querétaro through the “Química es evolución” program (Grant: In the process of being assigned).

## Conflicts of Interest Statement

The authors have no conflicts of interest to declare.

## Data availability statement

The data that support the findings of this study are available from the corresponding author upon reasonable request.

## Author contribution

All authors contributed to the study conception and design. Material preparation, data collection, and analysis were performed by Abigail Correa-Olivares, and Amanda de la Caridad Lahera Champagne. The first draft of the manuscript was written by Luis Alberto Madrigal-Perez, and all authors commented on earlier versions. All authors read and approved the final manuscript.

## Notes

### Competing Interest Statement

The authors have declared no competing interest.

